# SinaPlot: an enhanced chart for simple and truthful representation of single observations over multiple classes

**DOI:** 10.1101/028191

**Authors:** Nikos Sidiropoulos, Sina Hadi Sohi, Nicolas Rapin, Frederik Otzen Bagger

## Abstract

Recent developments in data driven science, in particular computational biology, have led scientists to integrate data from several sources, over diverse experimental procedures, or databases. This alone poses a major challenge in truthfully visualising data, especially when the amount of data points varies between classes.

To aid the presentation of datasets with differing sample size we have developed a new type of plot overcoming limitations of current standard visualization charts. Plots like bar charts, violin plots, strip charts or box-and-whiskers plots may provide visual information about mean/median, variance of the data, number of data points or density distribution of data; still, only a combination of these plots may provide all relevant information.

We have designed a new and simple plot inspired by the strip chart and the violin plot that operates by letting the normalized density of points restrict the jitter along the x-axis. The plot displays the same contour as a violin plot, but resembles a simple strip chart for small number of data points. In this way the plot conveys information of both the number of data points, the density distribution, outliers and data spread in a very simple, comprehensible and condensed format. The package for producing the plots is available for R through the CRAN network (https://cran.r-project.org/web/packages/sinaplot/index.html). In order to aid users without experience in R we also provide access to a web-server accepting excel sheets to produce the plots (http://servers.binf.ku.dk:8890/sinaplot/).

## Introduction

Visualization has been a key to gaining insights from data and conveying scientific results and messages in the most short, reliable and objective manner possible[1]. Recently, with the increasing amount of data available in all scientific disciplines, the task of visualizing data has become ever more important and difficult. Many attempts have been made to overcome the limitations of e.g. violin plots [2] or box-and-whiskers plots (attributed to [3] by [4]) by overlaying a jittered strip chart onto the plot.

This adds information on how outliers are spread, if there are any modality within a class, and gives a quick idea of whether some classes have more samples than others; but it also leaves a rather ink-intensive, and often quite messy-looking plot that can be difficult to read.

We have approached this problem by designing a plot that outlines the density, like the popular violin plot, while retaining all of the data points visible. This synergises the advantages of both the violin plot and the strip charts. The representation of the data as a whole remains simple, the density distribution is apparent, but the plot still provides information on how many data points are present in each class and if outliers are driving the tails of the distribution. In this way it is possible to convey information about the mean/median of the data, the variance of the data and number of data points or density distribution of data. The package should be useful for any scientist who wishes to visualize multiple observations of single variables over multiple classes.

## Methods

The SinaPlot estimates the kernel density for each class (e.g. all data points belonging to the same label group) and makes use of the density function to control of the jitter width. Specifically, we use the function density from package *stats* in *R* to estimate the density function of every class separately. The bandwidth can be adjusted to sharpen or smoothen the resulting curve. Subsequently, for each class, the range of the samples’ values is divided into bins of equal length. Samples within the same bin will be referred to as neighbours and the bin as neighbourhood. For neighbourhoods with more than 1 sample (value can be adjusted using the *neighbLimit* argument) the samples are spread along the x-axis by drawing values from a uniform distribution, setting the minimum and maximum values to the mean density estimate for the current neighbourhood. The process is illustrated in Figure 1.

**Figure 1:**
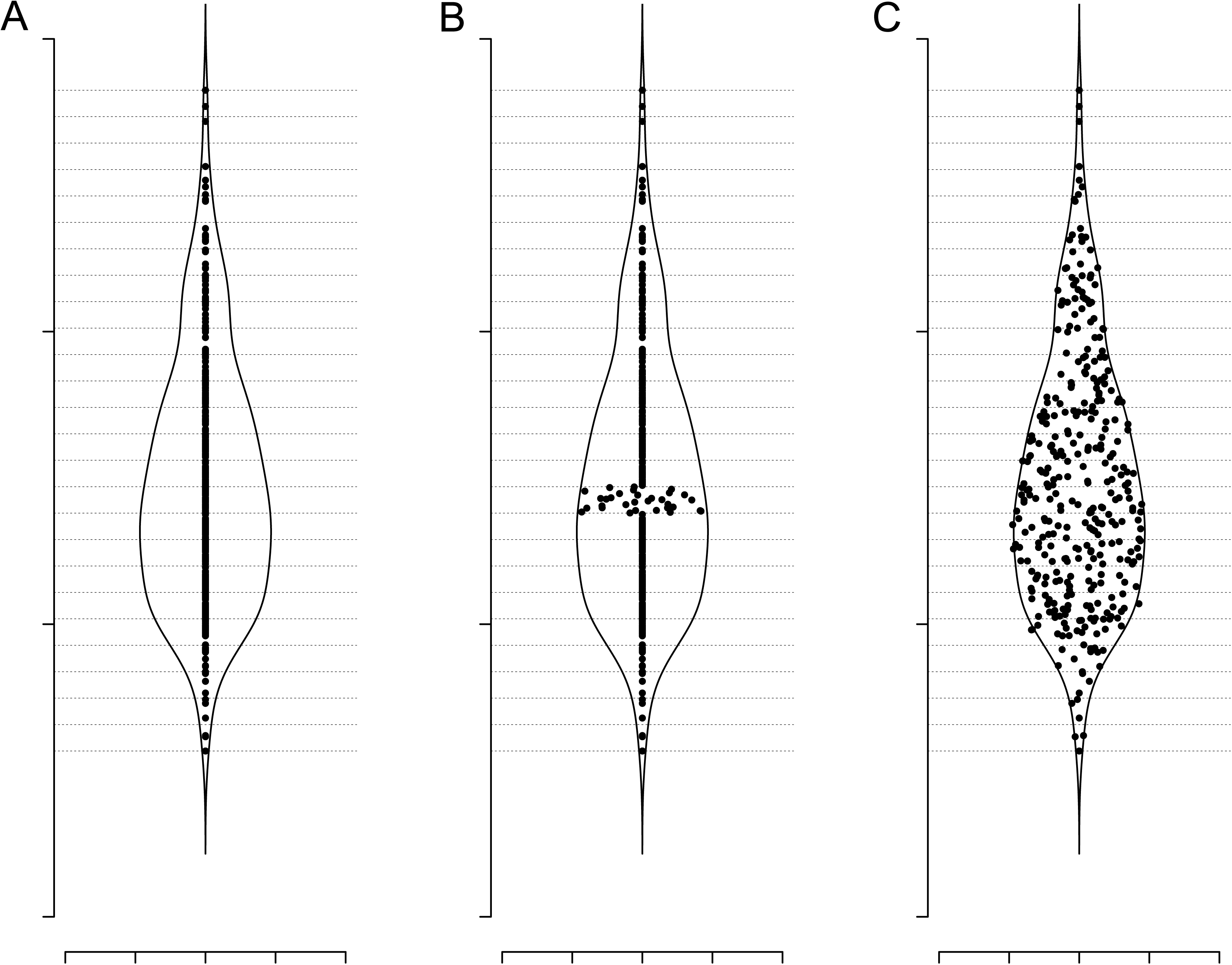
Jitter width estimation and spread. (A) Plot of a single class dataset overlayed by the samples’ density function along the y-axis. The range of the samples along the y-axis is divided into bins of equal length (horizontal dashed lines). (B) New x coordinates for samples that populate the same bin are drawn from a uniform distribution where the minimum and the maximum values correspond to the mean value of the density for that bin. (C) The same process is applied to all bins with sample size higher than 1 (value can be adjusted).

In the case of multiclass plotting the jitter width of each neighbourhood is scaled by default to represent a percentage of the neighbourhood with the highest sample density across all classes.

Alternatively, the jitter width can be estimated using the neighbour count within each class instead of the density curve. The widest borders are defined by the most highly populated neighbourhood and the rest neighbourhood borders are scaled based on it. The samples are spread within those borders by drawing their x-axis values from a uniform distribution as described above.

## Results

We have devised a new type of plot for visualization of large data-sets where multiple classes are represented (see Figure 2). The plot borrows from popular plot types for data visualization, such as beeswarm[5], boxplot, jitter plot, strip chart, and violin plot[2], and integrates their common advantages into a very simple and, most importantly, easily comprehensible chart. We have implemented an R-package[6] that can generate such plots to enable all data scientist to make use of this visualization method, and, furthermore we have devised a web-server for users with no experience with R. For a comparison with popular plots (see Figure 3), where the same data is plotted using 6 different plotting strategies. In contrast to standard usage of the violin plot the default setting in the SinaPlot is to set the width of the density distribution as normalized over all classes in the plot, and not within each class. This makes very differing numbers of data points quickly apparent. Hence, the widest distribution has the most points. This behavior can be set in the package by *scale* and on our web-interface.

**Figure 2:**
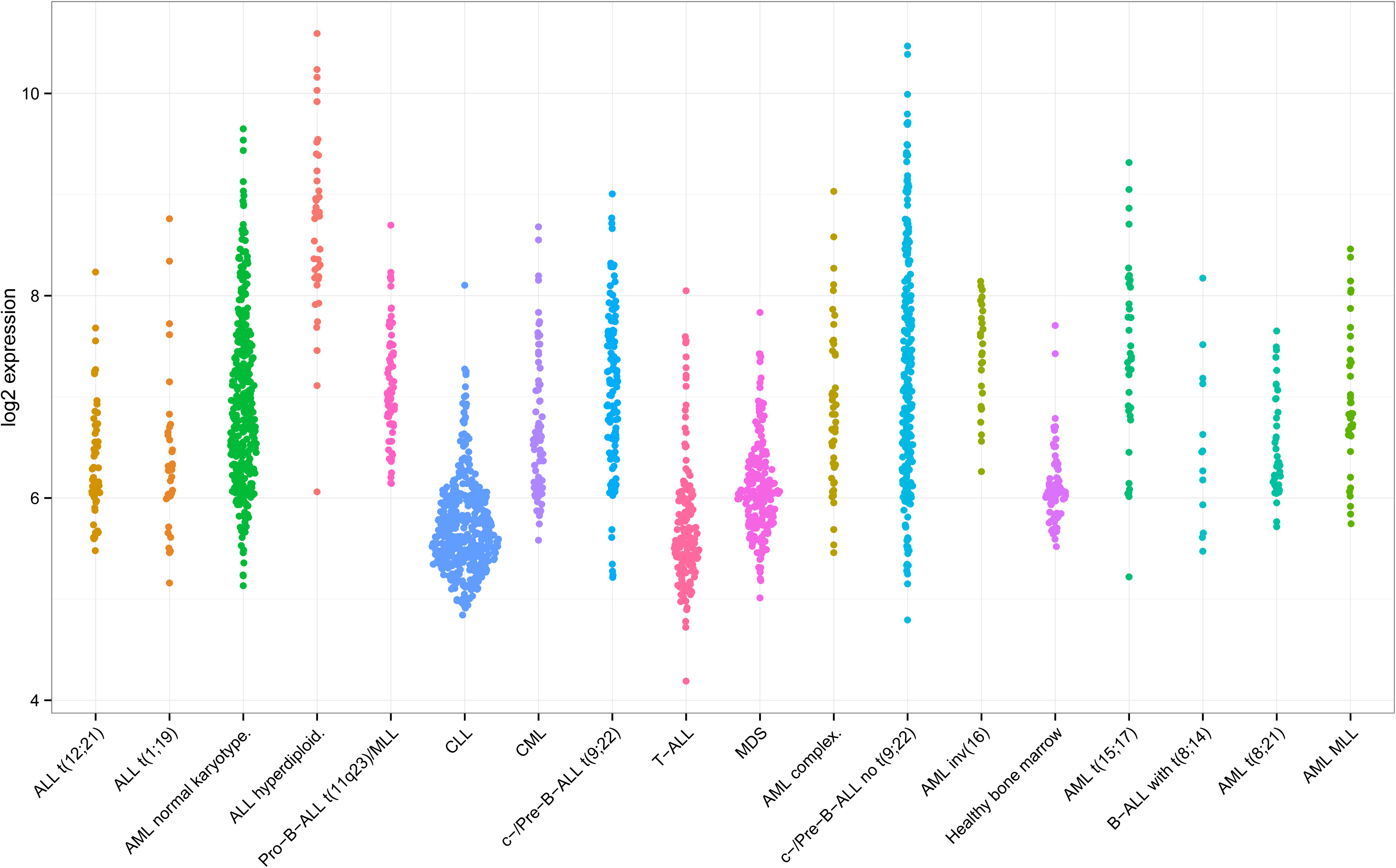
SinaPlot of gene IL3RA from a cohort of 2095 leukaemia patients obtained from [9] via (http://servers.binf.ku.dk/bloodspot).

**Figure 3:**
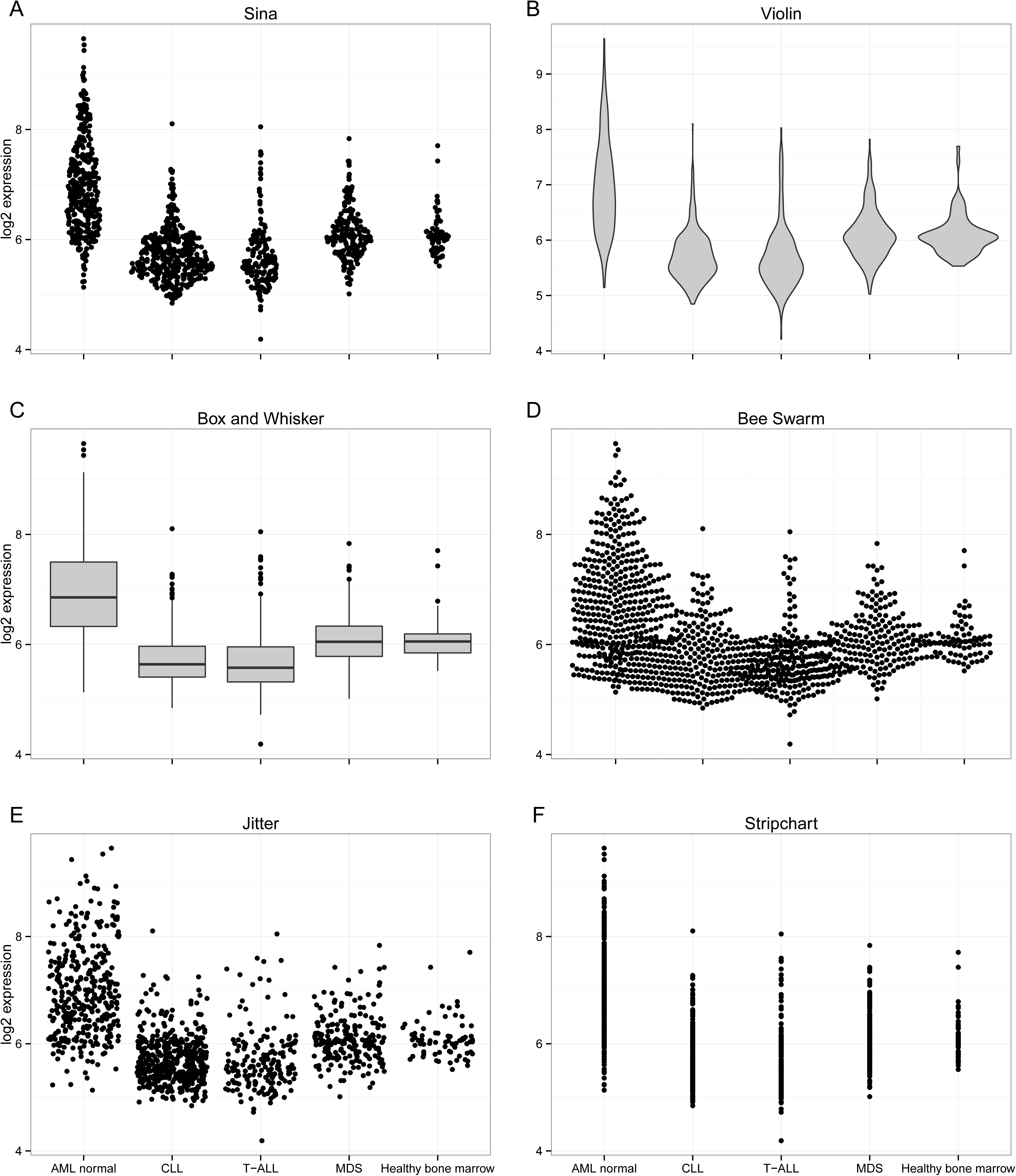
Expression values of gene IL3RA from 1252 AML/ALL and healthy bone marrow samples obtained from [9], grouped in 5 classes and visualised using SinaPlot (A) along with 5 other popular strategies; Violin plot (B), Box and Whiskers (C), Beeswarm (D), Jitter plot (E), Stripchart (F).

The R plotting function takes two mandatory arguments, *x* and *group* where *x* is a numeric vector of sample values to be split according to the grouping variable *group.* Several parameters can be tuned to control the y-axis bins, the density function and the sample scaling method, as well as to customize the graphical output.

The webserver is implemented as a shiny-R [6] web-server where the most important parameters are also available through a graphical interface. The server takes as input excel sheets, which we believe is a popular format for most scientist without programming experience.

We believe that any visualisation of multiclass data could benefit from using the SinaPlot, regardless of whether the number of observations in each category is equal over classes or differ extensively, as this fact is immediately apparent from the plot.

## Discussion

In the era of big data, ‘omics biology and integrative computer science, the need for simple and clear data visualization methods are becoming more and more important for understanding of underlying patterns in the data, especially when there are many observations or samples.

More and more studies rely on data that have been collected, produced or assembled for various unrelated purposes and by other teams of scientists. This, in effect, means that the number of measurements in each category is no longer controlled by the scientist, and it is very often not the same for each class. The same also goes for larger consortia data banks, where effort have been to truthfully represent the natural occurring incidence of a given disease phenotype, rather than collecting an equal sample size of each phenotype (TCGA is an example of such an effort[8]). With this type of large datasets, agglomerated representations have been used to avoid cluttering of the plots and convey messages about the underlying patterns to the target audience. We believe that there is a lack of intuitive understanding of how the data drives these plots, including box-and-whiskers plot and violin plots. Further, we find that circumventing this limitation by overlaying several plot types brings the problem of cluttered plots. We believe that the SinaPlot most truthfully conveys information in the data about number of samples, spread of the data, relation between the classes, outliers, and the overall density of the data, including where the centre of mass is, including information about the mean/median and possible modality. We suggest that the width of the density distribution is normalized over all classes in the plot, so that it becomes apparent how many samples are present in each class, which is in contrast to the present standard in violin plots. This behavior e.g. whether to normalise the density distribution over all the data, or within each category is optional in our R-package and on our web-interface. In summary, we believe that the SinaPlot most truthfully and simply represents multiclass plotting of single variables. This data visualization method should be of interest to anyone who works with analysing and presenting data. In particular we believe that the growing ‘omics community, often analysing large and collated datasets, will benefit from our visualization method. The R-function, documentation and vignette for producing the plot is freely available from CRAN (https://cran.r-project.org/web/packages/sinaplot/index.html) and plotting via web-interface (http://servers.binf.ku.dk:8890/sinaplot/).

